# Identifying seaweeds species of Chlorophyta, Phaeophyta and Rhodophyta using DNA barcodes

**DOI:** 10.1101/2020.08.30.274456

**Authors:** Narra Prasanthi, Chinnamani Prasannakumar, D. Annadurai, S. Mahendran, A. H. Mohammed Alshehri

## Abstract

Strengthening the DNA barcode database is important for a species level identification, which was lacking for seaweeds. We made an effort to collect and barcode seaweeds occurring along Southeast coast of India. We barcoded 31 seaweeds species belonging to 21 genera, 14 family, 12 order of 3 phyla (*viz.*, Chlorophyta, Phaeophyta and Rhodophyta). We found 10 species in 3 phyla and 2 genera (*Anthophycus* and *Chnoospora*) of Phaeophyta were barcoded for the first time. Uncorrected p-distance calculated using K2P, nucleotide diversity and Tajima’s test statistics reveals highest values among the species of Chlorophyta. Over all K2P distance was 0.36. The present study revealed the potentiality of rbcL gene sequences in identification of all 3 phyla of seaweeds. We also found that the present barcode reference libraries (GenBank and BOLD) were insufficient in seaweeds identification and more efforts were needed for strengthening local seaweed barcode library to benefit rapids developing field such as environmental DNA barcoding. We also show that the constructed barcode library could aid various industrial experts involved in seaweed bio-resource exploration and taxonomy/non-taxonomic researches involved in climate, agriculture and epigenetics research in precise seaweed identification. Since the rise of modern high-throughput sequencing technologies is significantly altering bio-monitoring applications and surveys, reference datasets such as ours will become essential in ecosystem’s health assessment and monitoring.

## 1. Introduction

Seaweeds are marine macroalgae that inhabit the littoral zone and are significant in terms of marine ecology and economics (Dhargalkar and Pereira, 2005). Seaweeds were taxonomically distributed in 3 major phyla; 1. Ochrophycea, commonly called as brown algae because of its xanthophyll pigment ‘fucoxanthin’, 2) Chlorophyta, commonly called as green algae because of dominant chlorophyll pigments ‘a’ and ‘b’, and minor xanthophyll pigments; and 3) Rhodophyta, commonly called as red algae because phycocyanin and phycoerythrin pigments (O’Sullivan et al., 2010). Globally more than 4000, 1500, and 900 species of Rhodophyta, Phaeophyta and Chlorophyta, respectively were documented (Dawes, 1998). Chlorophyta and Rhodophyta were dominant in the tropical and subtropical waters whereas Phaeophyta dominates cold temperate waters (Khan and Satam, 2003). Seaweeds plays a vital role in contribution of pharmacologically active compounds as 30% of marine derived pharmaceutical formulations were from Seaweeds (Blunt et al., 2007). In marine biodiversity point of view, dynamics of seaweed diversity are of major concerns as they constitute 40% of all marine invasive species documented so far (Schaffelke et al. 2006).

Identification of seaweeds based on morphological characters is difficult as most genera were known for its diverse morphotypes. For example; *Durvillaea antartica* in response to local hydrodynamic conditions forms distinct morphotype (Méndez et al., 2019). Also convergent evolution has simultaneously occurred in numerous distantly related macroalgae resulting in similar forms which complicates morphology based identification. For example; the uniqueness of kelp forests defined by its stiff stipes was the result of 5 separate evolutions (Starko et al., 2019).

DNA barcoding involves sequencing a gene fragment from precisely identified specimens to form a database and facilitate species identification (even by non-experts) simply by comparing the same gene sequences sequenced from unidentified specimens (Hebert et al., 2003, Mitchell, 2008). DNA barcoding has proved to be an efficient techniques for monitoring marine flora associated fauna (Manikantan et al., 2020; PrasannaKumar et al., 2020) and macro-algae (Kucera et al., 2008; Lee and Kim, 2015; Montes et al., 2017; Bartolo et al., 2020). The CBOL (Consortium for the Barcode of Life) has proposed rbcL (RuBisCO large subunit) and matK (maturase K) as DNA barcodes for plants (Hollingsworth et al., 2009). Absence of matK in green algae (excerpt Charophyte (Sanders et al., 2003)) made them inappropriate for barcoding Chlorophyta (*Caulerpa* sp.). Hence there is an urgency in evaluating a universal barcode gene for all 3 phyla (Phaeophyta, Rhodophyta and Chlorophyta) of seaweeds. rbcL barcodes has proven to be a potential barcode gene for identification of natural, aquacultured and processed seaweeds (Tan et al., 2012;Saunders and Moore, 2013; Gu et al., 2020). rbcL gene has also proven its ability in unmasking overlooked seaweed diversity (Saunders and McDevit, 2013), identification of cryptic invasive (Saunders, 2009; Geoffroy, 2012) and exotic (Mineur et al., 2012; Montes et al., 2017) species, besides discovering new seaweed species (Griffith et al., 2017). Documenting and accounting seaweed DNA barcodes for precise identification is important as seaweeds responds to variable water currents with flexible morphology and strengths (Sirison and Burnett, 2019).

A comprehensive reference genetic library of seaweed DNA barcodes, coupled with taxonomic and geo-referenced data is required for precise utilization of DNA barcoding technology (Bartolo et al., 2020). Since the significance of biotic indices in climate change studies are increasing (Brodie et al. 2017) and number of new seaweed taxa being discovered is on rise (De Clerck et al. 2013), creating a barcode library for extant seaweed species are the need of the hour as it would facilitate the identification of existing and discovery of new species. Such libraries could also facilitate precise identification of seaweeds for commercial, ecological and legislative purposes (Bartolo et al., 2020).

Strengthening the local barcode reference database is of paramount importance in using reference database for an accurate species identification (Hleap et al., 2020) which is currently lacking for seaweeds (Bartolo et al., 2020). Though peninsular India contains more than 860 seaweed species (Kannan and Thangaradjou, 2007; Jha et al., 2009; Ganesan et al., 2019; Mantri et al., 2019), DNA barcoding efforts are limited to sequencing one (Bast et al., 2014a, b; Bast et al., 2015; Bast et al., 2016a, b) or two species (Mahendran and Saravanan, 2014) at a time. For a first time we sampled and barcoded wide range of seaweed species belonging to all three phylum *viz*., Phaeophyta, Rhodophyta and Chlorophyta. The main objective of the study is to test the efficacy of DNA barcode reference libraries in precisely identifying Indian seaweeds and built a barcode reference library for Indian seaweeds.

## 2. Materials and methods

### 2.1. Sample collection and identification

Seaweeds were handpicked (including submerged rocks) in 5 different sampling stations along southeast coast of India during Jan, 2018 to Dec, 2019. Within each sampling stations, single to multiple sampling sites were chosen for collection. The 5 transects *viz*; Chennai (3 sites - Kovalam, Muttukadu & Mahabalipuram), Puducherry (3 sites – Marakkanam, Buckingham Canal bridge & Puducherry), Cuddalore (1 site - Parangipettai), Ramanathapuram (3 sites - Thondi, Mandapam & Rameshwaram), Tuticorin (1 site - Hare Island). Detailed geographical positions of the individual sampling sites were tabulated (Table 1). The seaweeds along with its holdfast were collected during low tide in the intertidal and sub-tidal region where the vegetation was discontinuous and occurring in patches. The samples were washed thoroughly with seawater to remove sand and other debris (marine soil debris, attached shell, mollusks adhering debris and associated biota). The second washing was done with deionised water to get rid of excess salts (Sivasankari et al., 2006). After washing, the samples were transported to the lab in sterile zip lock bags in cold conditions. The samples were kept in −20°C till further use.

**Table 1:** Geographical position of the collection site along southeast coast of India (Tamil Nadu coast) were represented in decimal degrees. Sampled locations were listed from northernmost (Chennai) to southernmost (Tuticorin) stations.

| S. No. | Transect | Sampling site | Latitude (N°) | Longitude (°E) |
| --- | --- | --- | --- | --- |
| 1 | Chennai | Muttukadu | 12.8265 | 80.2375 |
|  |  | Kovalam | 12.7840 | 80.2559 |
|  |  | Mahabalipuram | 12.6170 | 80.2001 |
| 2 | Puducherry | Marakkanam | 12.2046 | 79.9350 |
|  |  | Puducherry | 11.9074 | 79.8277 |
| 3 | Cuddalore | Parangipettai | 11.4941 | 79.7701 |
| 4 | Ramanathapuram | Thondi | 09.7483 | 79.0270 |
|  |  | Mandapam | 09.2870 | 79.1310 |
|  |  | Rameshwaram | 09.2962 | 79.2696 |
| 5 | Tuticorin | Hare Island | 08.7690 | 78.1776 |

The samples were thawed in artificial seawater in the lab. Morphology of all the Seaweed samples were carefully examined. The morphological and anatomical characteristics of the collected seaweeds were observed and identified using microscopic and macroscopic (such as frond size, leaf shape, leaf border, vesicles (air bladder), and receptacles) comparative analyses. The typical characteristics taken into account include internal structure, color, size & shape and by comparing to the existing photographs and data previously published for this region (Dinabandhu, 2010). The seaweeds were also identified based on the taxonomic keys described by Srinivasan (1969) & (1973), Rao (1987), Chennubhotla et al., (1987), Ganesapandian and Kumaraguru (2008), Jha et al., (2009) and with catalogue of benthic marine algae of Indian Ocean (Silva et al., 1996). Total of 31 species were identified and the voucher specimens were named from (DNA Barcoding of Indian Seaweeds) DBIS1 to DBIS31. The same name tags were used for molecular analysis. The metadata including pictures and systematic positions of the identified species could be accessed in Barcode of Life Database (BOLD; www.boldsystems.org) under the project title “DNA barcoding of Indian Seaweeds” or using a unique tag “DBISW”.

### 2.2. DNA isolation, PCR and sequencing

Total genomic DNA was extracted using HiPurA™ marine Algal DNA extraction kit (HiMedia Labs. Pvt. Ltd., India) as per manufacturers’ instructions and confirmed in 1.2% agarose gel. 100μl of elution buffer provided with the kit was used as negative control in DNA extraction. The 20μL PCR reaction mix contained 2μl of 10X reaction buffer with 15mM MgCl_2_ (Applied Biosystems, India), 4μl each of 10mM primer, 2μl of 1mM dNTPs (Bioserve Biotechnologies, India), 0.5μl of Dream-Taq DNA polymerase 0.5 unit) (Bioserve Biotechnologies, India), 4μl of template DNA and 7.5μl of double distilled water. The target rbcL gene region was amplified using universal plant DNA barcoding primers (CBOL, 2009), rbcLa-F: 5’-ATGTCACCACAAACAGAGACTAAAGC-3’ and rbcLa-R: 5’-GTAAAATCAAGTCCACCRCG-3’. PCR was performed using a reaction mixture of a total volume of 25μl; 12.5μl of Taq PCR Master Mix (Invitrogen, India), 11μl distilled water, 0.5μl forward primer (10 μM), 0.5μl reverse primer (10 μM), and 0.5μl of the DNA template (50– ng/μl). The PCR conditions were as follows: 1 cycle (94 °C for 3 min), 35 cycles (94 °C for 1 min, 55 °C for 1 min, and 72 °C for 1 min), and 1 cycle 72 °C for 7 min. The same negative control used during DNA extraction acted as negative control for PCR. PCR products of rbcL gene fragment was checked on 1.5% agarose gel electrophoresis (using above protocol described in 5.2.2.) for 600bps amplicon. DNA sequencing was performed at Macrogen Inc. (Seoul, South Korea).

### 2.3. Sequence analysis

For few samples, PCR amplification and DNA sequencing reactions were repeated either to improve the length of DNA sequences recovered or the quality of the final chromatograph. Only good quality sequences (with precise base calling and 90% of total length) were included in the study. All sequence chromatographs were manually double checked for quality using Chromas Pro version 2.6.6. Forward and reverse chromatograms were aligned using Bio Edit (Hall, 1999). The sequences were aligned in Clustal X ver. 2.0.6 (Thompson et al., 1997) and Molecular Evolutionary Genetic analysis (MEGA) ver. X was used for phylogenetic and pair-wise distance analysis (Kumar et al., 2018). The pair-wise distance was calculated as per Kimura-2 parametric distance model (Kimura, 1980). NJ tree was redrawn using Interactive Tree Of Life (iTOL) (Letunic and Bork, 2019) for better representation of tree based identification. Kimura-2 parameter distance model (Kimura, 1980) was to calculate the distances between the sequences. All three codon positions and non-codon positions were included and all the alignment positions containing gaps and missing data was eliminated from the analysis. MEGA X was also used to conduct nucleotide diversity and Tajima’s neutrality (Tajima, 1989; Nei and Kumar, 2000) tests. The rbcL sequences produced in the present study was available for public in Genbank and could be accessed through accession numbers MT478065-MT478095 and in BOLD through http://dx.doi.org/10.5883/DS-DBISW.

## 3. Result and discussion

### 3.1. Species composition and BLAST analysis

DNA from 31 samples were successfully extracted and PCR amplified, with no extracted DNA or PCR amplification in negative controls. Among 31 species collected, Rhodophyta, Phaeophyta and Chlorophyta contributed 14, 9 and 8 species, respectively. Among 31 seaweeds species barcoded, there were 21 genera belonging to 14 family, 12 order, 3 classes (viz., Florideophyceae, Phaeophyceae and Ulvophyceae) in 3 different phyla (viz., Chlorophyta, Phaeophyta and Rhodophyta). List of species barcoded with its species authority and corresponding GenBank accession number is given in table 2.

**Table 2:**
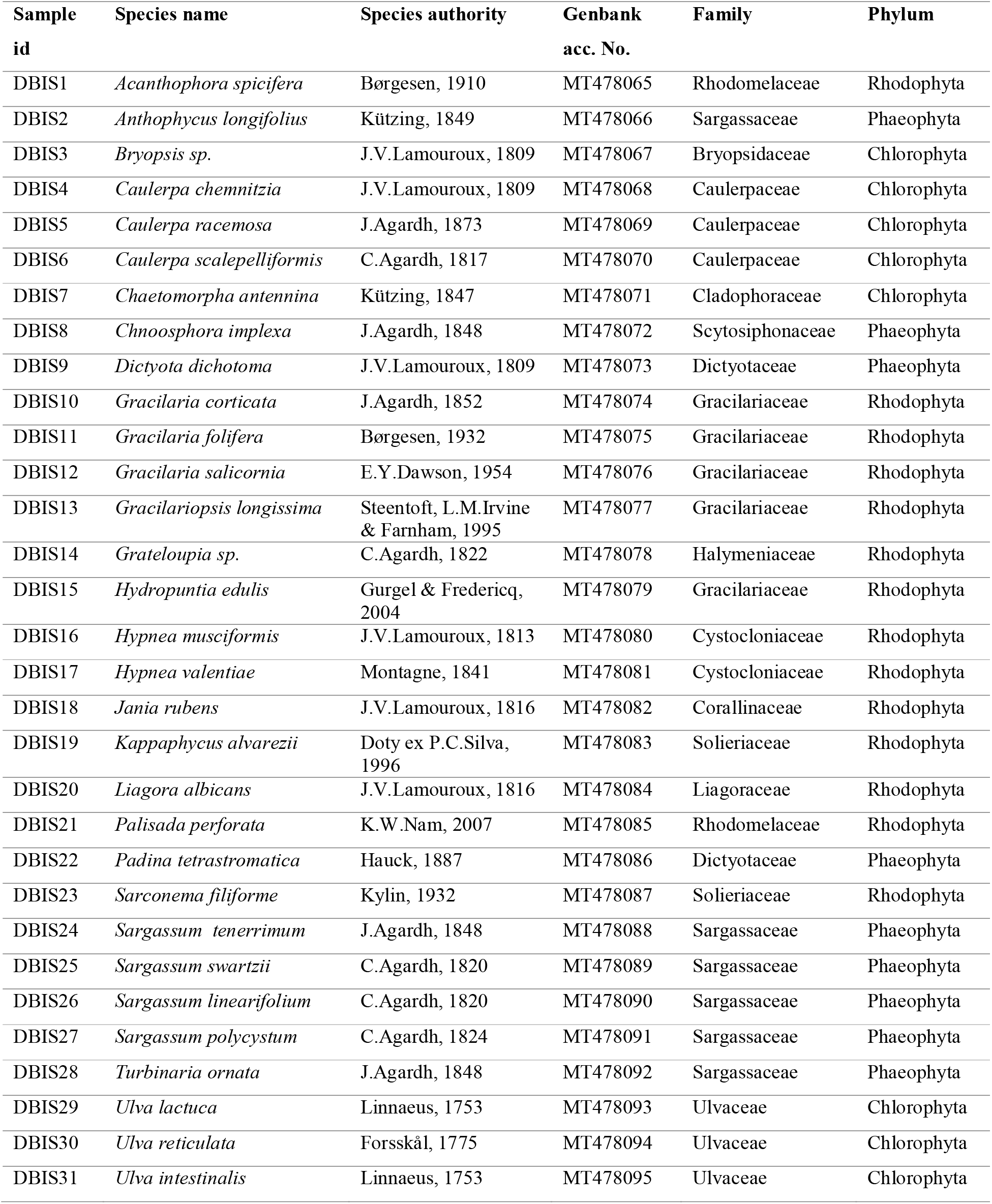
List of species collected and barcoded with its corresponding GenBank accession numbers and sample id.

Among 31 species barcoded, BLAST analysis could precisely identify 61.3% (n=19 species) of sequences with maximum of 99.33% similarity (for *Ulva lactuca*) (Table 3). We found that 38.7% (n=12 species) of rbcL sequences was barcoded for the first time ever, as the corresponding sequences were matched with less similarity with other genera and the individual search of barcoded species in Genabank showed its absence. Among 12 species sequenced for its rbcL region, none of the members of 2 genera (phyla: Phaeophyta) *viz*; *Anthophycus* sp. and *Chnoospora* sp. was present in the Genbank, which means through submission of rbcL sequences (of *A. longifolius* and *C. implexa*), those two respective genera was represented for the first time in Genbank. Apart from *A. longifolius* and *C. implexa*, other members of Phaeophyta barcoded for the first time were, *Sargassum tenerrimum, S. swartzii, S. linearifolium*. Among the 12 species barcoded for the first time, 3 members of Rhodophyta were *Gracilaria corticata, G. folifera, Grateloupia* sp. and 4 members of Chlorophyta were *Caulerpa chemnitzia, C. scalpelliformis, Chaetomorpha antennina, Ulva intestinalis*.

**Table 3:**
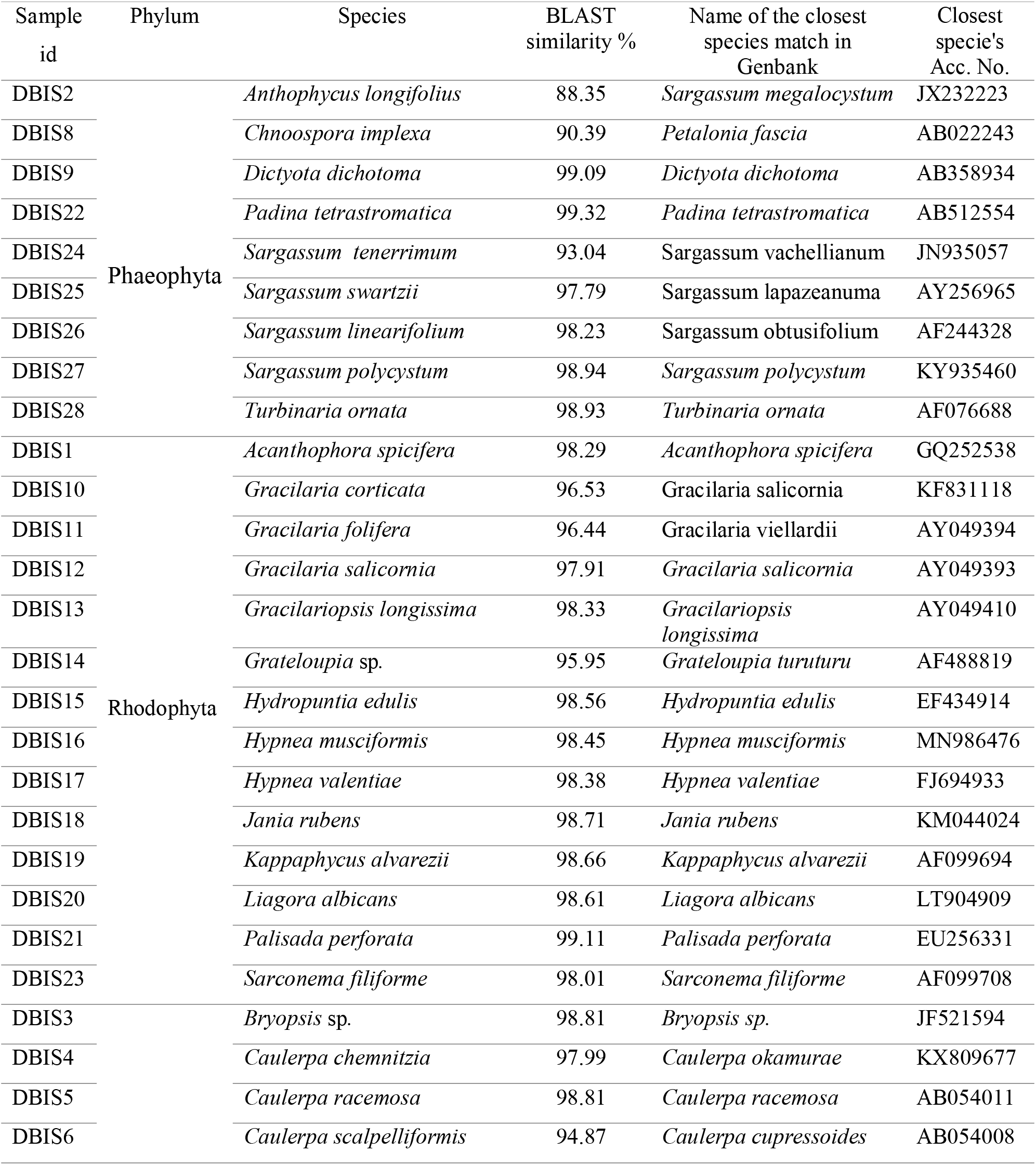
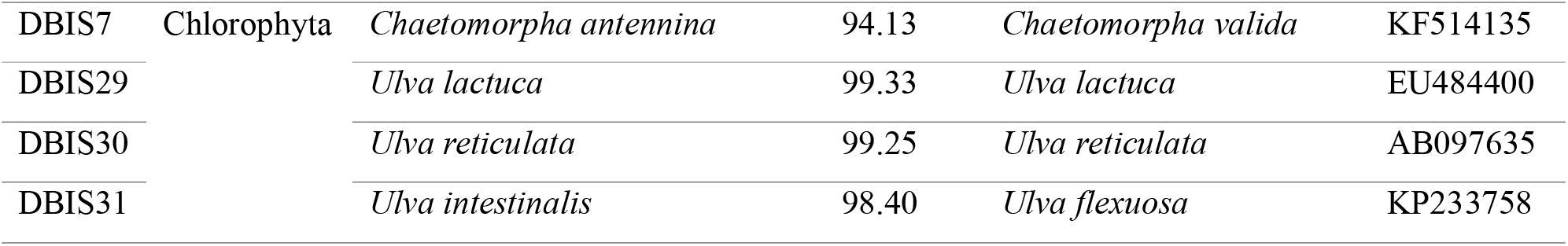
BLAST analysis of rbcL sequences produced in this study against the GenBank reference sequences with the name of the closest match and corresponding accession number.

DNA barcode match results given by BOLD search engine, consisting of score summary with top 99 matches and e-value summary could be assessed through https://www.doi.org/10.13140/RG.2.2.12833.22889

### 3.2. DNA barcoding Phaeophyta and its utility

Among the 9 species of Phaeophyta barcoded in the present study, *Chnoospora implexa* are well known for its anti-microbial properties (Seker et al., 2019; Rani et al., 2020). Whereas *Dictyota dichotoma* are known for hosting rich mycobiota diversity (Pasqualetti et al., 2020) and its anti-cancer potential (El-Shaibany et al., 2020). In recent years, genetic characterization of *D. dichotoma* under acidified ocean conditions receives special attention (Porzio et al., 2020). *Padina tetrastromatica* and *P*. *tetrastromatica* barcoded from the same sampling location were known for high mineral compositions (Vasuki et al., 2020) and antiviral (HIV) activities (Subramaniam et al., 2020) respectively. Hence generating barcodes for the above said species would benefit help non-taxonomists experts from mineral and pharmaceutical industries for species identification. *Turbinaria* spp. were known to contain diverse pharmacologically active formulations (Rushdi et al., 2020), extracts of *T. ornata* barcoded in the present study was known to synthesise silver nano-particle (Renuka et al., 2020) and nano material (Govindaraju et al., 2020). DNA barcoding *T. ornata* is significant as they exhibit dynamic morphological characteristics based on the strength of ambient water currents (Sirison and Burnett, 2019).

*Sargassum* spp. known for delivering wide range of natural and pharmaceutical formulations (Rushdi et al., 2020), significantly influences the coastal ecosystems (Nguyen and Boo, 2020) and the species barcoded in the present study such as *S. polycystum* forms a thick mat like growth which profoundly influences the coastal waters (example, Vietnam coast (Nguyen and Boo, 2020)). *S. tenerrimum* barcoded in the present study was previously known for its application as catalyst in biochar production (Kumar et al., 2020) and in lead absorption (Tukarambai and Venakateswarlu, 2020). DNA barcoding the *S. swartzii* which was estimated to play important role in studying the future ocean temperature rise (Graba-Landry et al., 2020) will facilitate easy identification for climate researchers. DNA barcoding the seaweed species such as *Sargassum linearifolium* which forms its own ecosystems by hosting diverse invertebrate communities (Lanham et al., 2015) and exhibit with high intraspecies variability and adaptive morphological changes (Stelling□Wood et al., 2020) gains more importance. *Sargassum polycystum* barcoded from the present sampled area were also known for its wide range of phytochemical composition (Murugaiyan, 2020). Hence the Phaeophyta barcodes produced in the present study will be useful to taxonomic, non-taxonomic experts, pharmaceutical, minerals, naturopathic and nano-technology industries, researchers of climate science, agriculture and epigenetics.

### 3.3. DNA barcoding Rhodophyta and its utility

*Acanthophora spicifera* barcoded in the present study are known for branched-long cylindrical thallus (Nassar, 2012) and preferred habitat for brachyuran (Granado et al., 2020) and horse shoe crab species (Butler et al., 2020) whose occurrences is seasonal (example; South-eastern Brazil coast (Lula Leite et al., 2020)). *A. spicifera* sampled from present sampling study area was also known to accumulate Cadmium in its tissues which were biomagnified to its animal in-habitants (Ganesan et al., 2020). Hence generating DNA barcodes for such species will aid in effective environmental monitoring. *Gracilaria corticata* barcoded from the present sampling area were known to for its high sulphate and mineral content, which was currently utilized by food industry (Subramanian et al., 2020). *G. cortica* extracts plays vital role in Zinc removal (Heidari et al., 2020), Zinc oxide (Nasab et al., 2020) and silver nano-particle synthesis (Rajivgandhi et al., 2020). The extracts of *Gracilaria folifera* barcoded in the present study were known for mosquito larvicidal activities (Bibi et al., 2020).

*Gracilaria salicornia* barcoded in the present study was an invasive species of Hawaiian coastal waters (Hamel and Smith, 2020) and less preferred substratum for microbial grazing (Tan et al., 2020). It was also known that when *G. salicornia* and *Acanthophora spicifera* co-occurs, epiphytic micro-algal assemblages prefers *A. spicifera*, rather than *G. salicornia* (Beringuela et al., 2020). Hence the DNA barcodes of *G. salicornia* generated in the present study will be useful for exploring invasive potential and in macro-algal identification during microbial-macro-algal interaction studies. *Gracilariopsis longissima* are actively cultured seafood (Bermejo et al., 2020) and a source of bioactive compounds (Susanto et al., 2019). *Hydropuntia edulis* were known for its high content of UV-absorbing compounds (Tanaka et al., 2020). Hence the generated barcodes will be useful for taxonomic non-experts of food, pharmaceutical and cosmetic industry. However it’s worth mentioning that the density of *H. edulis* in the present sampled area was alarmingly declining due to un-sustainable harvesting usually for local food grade agar production (Rao et al., 2006; Ganesan et al., 2011).

*Hypnea musciformis* barcoded in the present study were known for bio-preservative compounds which improves shelf life period of seafood (Arulkumar et al., 2020). Previous studies has shown that biochemical composition (protein, carbohydrate and lipids) of *Hypnea valentiae* were known to vary seasonally from the sampled area (Murugaiyan and Sivakumar, 2020). Further studies could be carried out to explore the possibility of linking DNA barcodes to inter and intra-species biochemical constituents of seaweeds. *Jania rubens* that hosts diverse amphipod species (Kh. Gabr et al., 2020) are also known for its diverse haplotypes (Harvey et al., 2020) and bioactive resources (Rashad and El-Chaghaby, 2020). Further studies could explore, if DNA barcodes could effectively delineate various haplotypes of *J. rubens*. In species like *Kappaphycus alvarezii* barcoded in the present study and the ones occurring in Brazilian coast (Nogueira et al., 2020), various haplotypes were known to contain variable composition of anti-oxidants (Araújo et al., 2020). They were also actively used for the nutrient removal in integrated fish culture systems (Kambey et al., 2020). DNA barcodes of *Palisada perforata* produced in the present study can also be used to identify the same species occurring in Egyptian (Kh. Gabr et al., 2020) and Persian coasts (Abdollahi et al., 2020), as DNA barcoding works universal. The carrageenans of *Sarconema filiforme* barcoded in the present study were known for anti-inflammatory and prebiotic activity (du Preez et al., 2020), which was an anthropogenically threatened species in Tanzanian coast (Kayombo et al., 2020). Hence the generated barcodes will aid in environmental monitoring for ensuring continuous perpetuation of this species. DNA barcodes of Rhodophyta generated will be useful for species identification by non-experts of seafood (and its by-prodcuts), pharmaceutical, cosmetic and nano-technological industry and in environmental monitoring to explore the presence of invasive and optimal perpetuation of threatened seaweed species besides the morphological variability exhibited by Rhodophyta (Rodríguez and Otaíza, 2020).

### 3.4. DNA barcoding Chlorophyta

DNA barcodes of *Caulerpa chemnitzia* could be useful for researchers of phytochemical industries. Phytochemical and bactericidal activity of *C. chemnitzia* barcoded in present study area were known for high quantities of terpenoids, tannins and phenolic resins (Krishnamoorthy et al., 2015). *C. chemnitzia* were also known to occur in Bangladesh coastal waters (Bay of Bengal) (Abdullah et al., 2020). Recently sulfated polysaccharide of *Caulerpa racemosa* were positively evaluated for anti-inflamatory activities (Ribeiro et al., 2020). The extracts *C. racemosa* were also used as sun screens (Ersalina et al., 2020) and source of anti-bacterial compounds (Belkacemi et al., 2020). *C. racemosa* collected from same sampled are of the present study were also known for bio-diesel production (Balu et al., 2020). Hence the DNA barcodes of *C. racemosa* will be useful for researchers of pharmaceutical, cosmetic and bio-diesel industries for seaweed species identification. The edible species, *Caulerpa scalpelliformis* barcoded from same sampled area were also known to bio-accumulate various heavy metals (Rajaram et al., 2020), increasing the chances for exploring this species in bio-remediation applications.

*Chaetomorpha antennina* barcoded in the present study were known for its fastidious growth patterns by rapid nutrient uptake (Imchen and Ezaz, 2018). Correlating DNA barcodes with ambient seawater conditions could aid in finding optimum species for the given ecosystems in near future. *Ulva lactuca* barcoded in the present study were good bio indicator of trace metal contamination (Bonanno et al., 2020), which are known for rapid absorption of cadmium metals (El-Sheekh et al., 2020). Recently they are used as model organisms for energy budget studies for managing diverse ecological conditions (Lavaud et al., 2020). Hence the DNA barcodes of *U. lactuca* would of immense use in environmental monitoring. *Ulva reticulata* barcoded in the present study was known for high carbon sequestration potential (Sathakit et al., 2020) and its biomass were used for bio-diesel and bio-ethanol production (Osman et al., 2020).

### 3.5. Tree based identification

NJ tree precisely clustered members of 3 different phyla into 3 different groups (Fig. 1). Among the members of Phaeophyta, the rbcL sequences of *Dictyota dichotoma, Padina tetrastromatica, Turbinaria ornate*, and *Sargassum polycystum* retrieved from Genbank precisely matched with the same members sequenced in the present study. Since other rbcL sequences belonging to *Anthophycus longifolius, Chnoosphora implexa, Sargassum tenerrimum, S. swartzii*, and *S. linearifolium* was sequenced for the first time, no representative match could be found for tree based identification. Among the members of Chlorophyta, rbcL reference sequences of *Bryopsis* sp., *Caulerpa racemosa, Ulva lactuca, Ulva reticulate* precisely grouped with the same species sequenced in the present study whereas the species barcoded for the first time *viz*; *Caulerpa chemnitzia, C. scalpelliformis*, *Chaetomorpha antennina* and *Ulva intestinalis* did not have exact match in Genbank. The phylogenetic relaionships observed between *Caulerpa racemosa* and *C. scalpelliformis* was similar to that of previous studies (Kazi et al., 2013). Also phylogenetic relationship observed between *Caulerpa racemosa* and *Ulva reticulata* was similar of those observed in the previous study (Mahendran and Saravanan, 2017).

**Fig. 1 :**
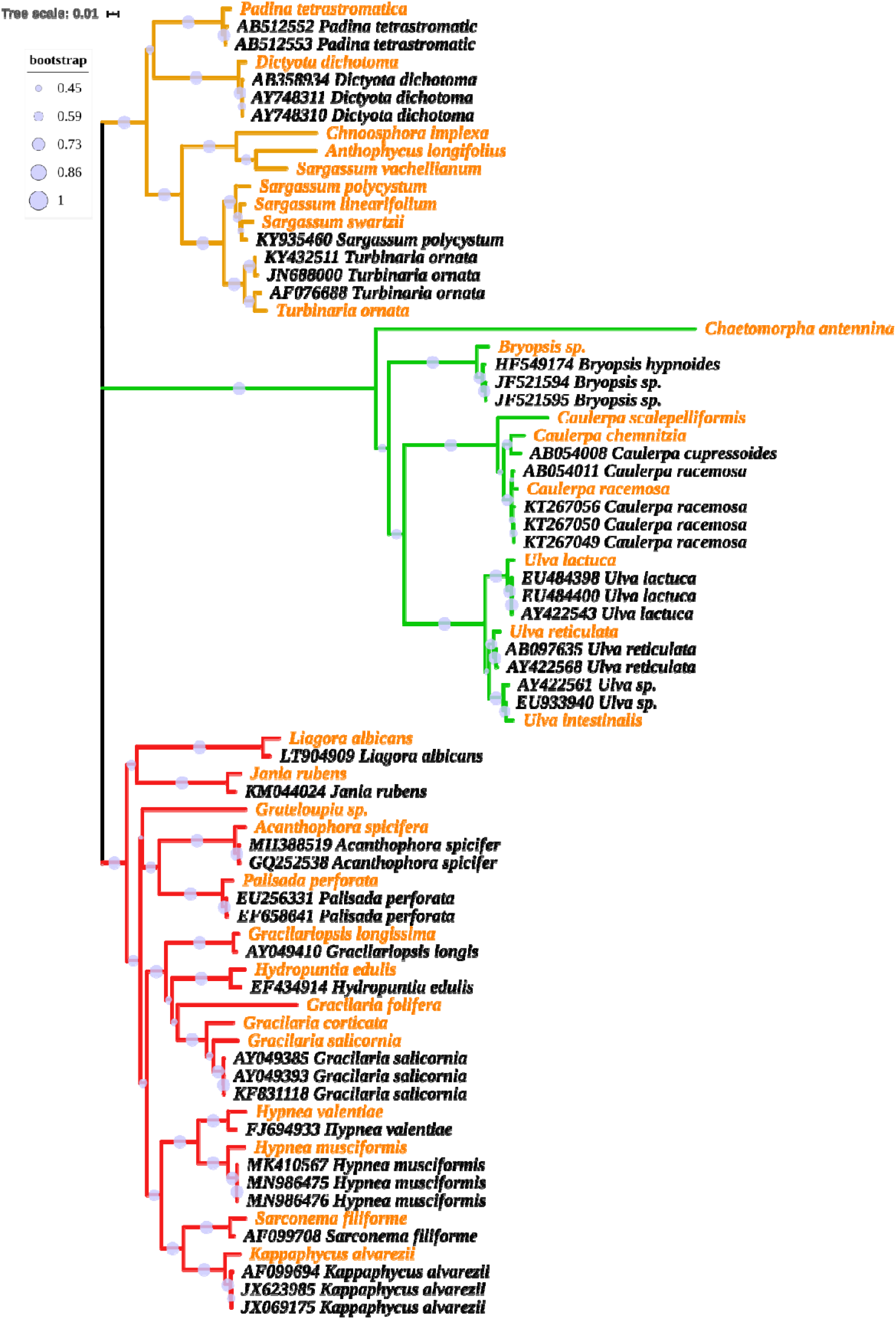
NJ tree drawn using rbcL sequences of 31 seaweeds species collected in the present study using Kimura-2 parameter model. Members of Phaeophyta, Chlorophyta and Rhodophyta are grouped in orange, green and red clades respectively. Sequences retrieved from GenBank were indicated by “accession-number_species-name” in the tree. Example, “JX069175 *Kappaphycus alvarezii*”, whereas the sequences generated from the present study was indicated without accession numbers and are distinguished by orange colour fonts.

Among the member of Rhodophyta, the reference sequences (of *Acanthophora spicifera, Gracilaria salicornia, Gracilariopsis longissima, Hydropuntia edulis Hypnea musciformis*, *Hypnea valentiae, Jania rubens, Kappaphycus alvarezii, Liagora albicans, Palisada perforate, Sarconema filiforme*) precisely grouped with same species sequenced in the present study, whereas the first time barcoded species viz; *Gracilaria corticata, Gracilaria folifera, Grateloupia* sp. did not have significant match in the Genbank.

Though previous study identified rbcL as potential candidate for species delineation in Chlorophyta (Kazi et al., 2013) and not as potential as mitochondrial genes in Rhodophyta (Ale et al., 2020; Siddiqui et al., 2020), the present study through tree-based identification reveals that rbcL as a potential candidate for barcoding seaweeds belonging to all 3 phyla. Alshehri et al. (2019) with limited sampling of seaweeds (n= 8 species) from all 3 phyla has also shown that rbcL could aid in precise species identification.

### 3.4. Genetic distance analysis

Kimura 2-parametric (K2P) (or uncorrected p) distance in all three phyla was positively correlated with nucleotide diversity (π) and Tajima’s statistics (D). K2P, π and D was higher in Chlorophyta (0.23, 0.2 and 1.1) followed by Rhodophyta (0.2, 0.17 and 1.15) and Phaeophyta (0.15, 0.13 and 1) respectively (fig. 2). Kazi et al. (2013) found the overall K2P distances among the *Caulerpa* spp. (Chlorophyta) was 0.043 which is well within the distance value of Cholorophyta (0.23) (derived using rbcL sequences belonging to 8 species, including 3 *Caulerpa* spp.) in the present study. Among the Rhotophyta members barcodesd previous (Tan et al., 2012), overall K2P distances of 0.06 was observed which is well within 0.19 observed in the present study. It is evident that better picture of average K2P distances among various seaweed phyla could be obtained with increase species coverage. Over all K2P distance, nucleotide diversity and Tajima’s test statistics for the 31 seaweed species were 0.36, 0.27 and 2.65, respectively.

**Fig. 2 :**
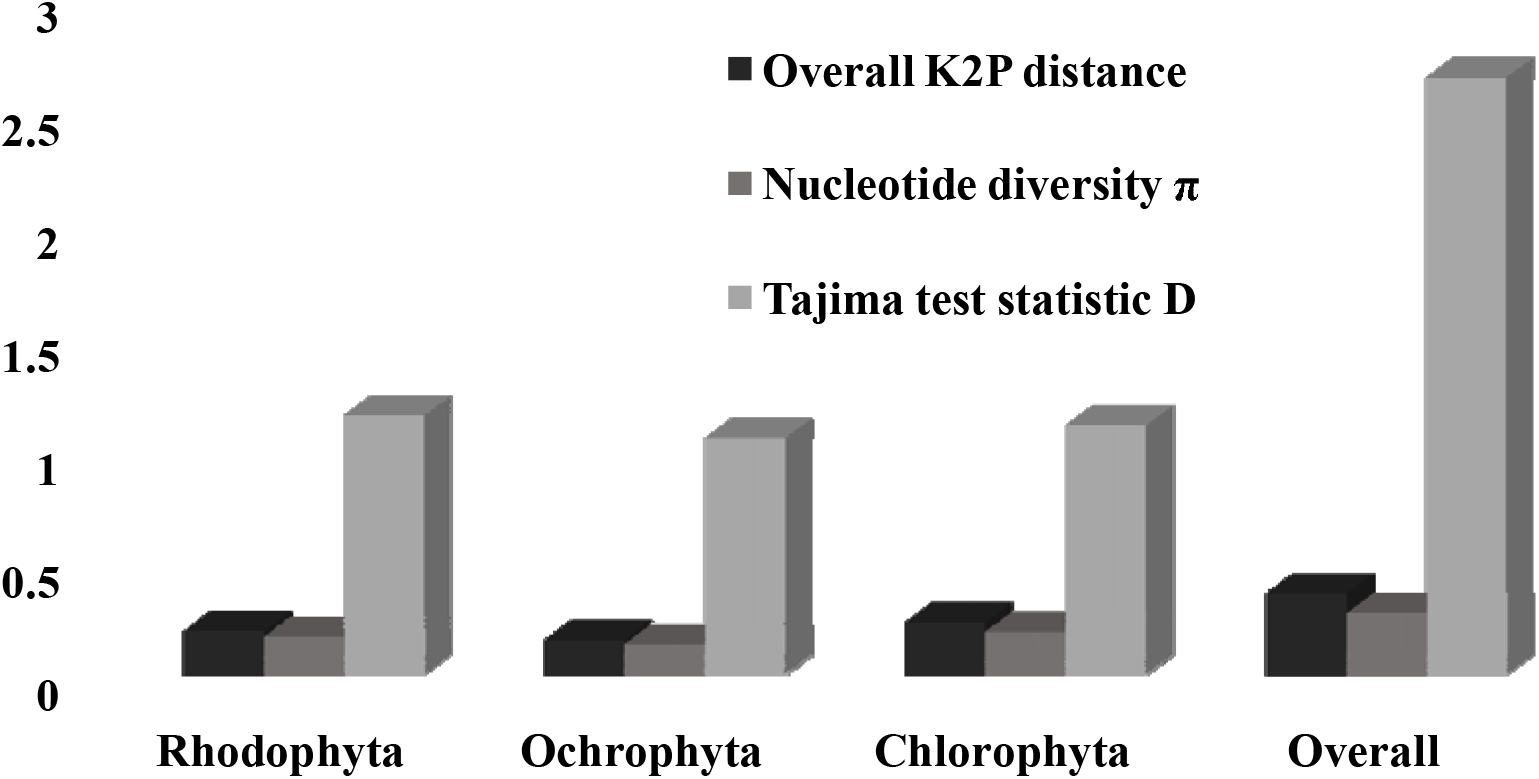
The graph shows the K2P distance, nucleotide diversity and Tajima’s test statistics for the seaweeds species of individual phyla and overall average. The values were in the trends of Average of 3 phyla (overall) > Chlorophyta > Rhodophyta > Phaeophyta.

## 4. Conclusion

We have synthesised a comprehensive barcode data for 31 seaweed species (with 12 species barcoded for the first time) occurring along the Southeast coast of India. The present barcode reference libraries are insufficient in marine macro-algal identification for Indian species and more efforts for DNA barcoding the local species is necessary to facilitate the environmental monitoring efforts. Building a comprehensive local barcode reference library could contribute to resolving macroalgal taxonomy and systematics and address biogeography pertaining to invasion of non-indigenous species. This could also result in the development and application of cost-effective and better biodiversity monitoring projects which could contributes to the EU Directives and UN conventions. Hence strengthening the local barcode libraries by barcoding all species (>800 Indian seaweed species were documented globally) could facilitate cost-effective biodiversity surveys and effective environmental barcoding programmes in near future. The generated barcodes will be useful for various industrial (Pharmaceutical, fuel, seafood, cosmetic and nano-technological) and research (climate change, species distribution) applications. However regular monitoring of seaweeds in the marine environment will be necessary to evaluate the ecological shifts due to climate change (Bringloe et al., 2019). The rise of modern high-throughput sequencing technologies will significantly alter bio-monitoring applications and surveys in the near future (Fonseca et al., 2010; Hajibabaei et al., 2011; Leray et al., 2015). As a result, reference datasets such as ours will become essential for assessing health and monitoring various aquatic environments using seaweed barcodes. Further studies could increase number of barcodes per species from same and different geographical locations to shed lights on phylogeographic signals to trace back the origin of drifting seaweeds (Guillemin et al., 2020).

## Acknowledgement

First authors are thankful to director, Centre of Advanced Study in Marine Biology, Annamalai University for the encouragement and facility provided. Corresponding author thank the Department of Science and Technology, Ministry of Science and Technology (GoI) for the INSPIRE fellowship (IF10431).

## Notes

https://www.doi.org/10.13140/RG.2.2.12833.22889

http://dx.doi.org/10.5883/DS-DBISW

### Competing Interest Statement

The authors have declared no competing interest.

### Summary of Updates

Latest references have been included

## References

Abdollahi, R., Ebrahimnezhad Darzi, S., Rahimian, H., & Naderloo, R. (2020). Biodiversity of intertidal and shallow subtidal habitats in Abu Musa Island, Persian Gulf, Iran, with an updated species checklist. Regional Studies in Marine Science, 33, 100975. https://doi.org/10.1016/j.rsma.2019.100975

Abdullah Al, M., Akhtar, A., Rahman, M. F., Kamal, A. H. M., Karim, N. U., & Hassan, Md. L. (2020). Habitat structure and diversity patterns of seaweeds in the coastal waters of Saint Martin’s Island, Bay of Bengal, Bangladesh. Regional Studies in Marine Science, 33, 100959. https://doi.org/10.1016/j.rsma.2019.100959

Ale, M., Barrett, K., Addico, G., Rhein-Knudsen, N., deGraft-Johnson, A., & Meyer, A. (2016). DNA-Based Identification and Chemical Characteristics of Hypnea musciformis from Coastal Sites in Ghana. Diversity, 8(4), 14. https://doi.org/10.3390/d8020014

Ali Alshehri, M., Aziz, A. T., Alzahrani, O., Alasmari, A., Ibrahim, S., Osman, G., & Bahattab, O. (2019). DNA-barcoding and Species Identification for some Saudi Arabia Seaweeds using rbcL Gene. Journal of Pure and Applied Microbiology, 13(4), 2035–2044. https://doi.org/10.22207/jpam.13.4.15

Araújo, P. G., Nardelli, A. E., Fujii, M. T., & Chow, F. (2020). Antioxidant properties of different strains of Kappaphycus alvarezii (Rhodophyta) farmed on the Brazilian coast. Phycologia, 59(3), 272–279. https://doi.org/10.1080/00318884.2020.1736878

Arulkumar, A., Satheeshkumar, K., Paramasivam, S., Rameshthangam, P., & Miranda, J. M. (2020). Chemical Biopreservative Effects of Red Seaweed on the Shelf Life of Black Tiger Shrimp (Penaeus monodon). Foods, 9(5), 634. https://doi.org/10.3390/foods9050634

Balu M, Lingaduraia K, Shanmugam P, Raja K, Bhanu Teja N and Vijayan V. (2020) Biodiesel production from Caulerpa racemosa (macroalgae) oil. Indian Journal of Geo Marine Sciences. 49(04): 616–621.

Bartolo, A. G., Zammit, G., Peters, A. F., & Küpper, F. C. (2020). The current state of DNA barcoding of macroalgae in the Mediterranean Sea: presently lacking but urgently required. Botanica Marina, 63(3), 253–272. https://doi.org/10.1515/bot-2019-0041

Bast, F., Bhushan, S., & John, A. A. (2014b). DNA barcoding of a new record of epi-endophytic green algae Ulvella leptochaete (Ulvellaceae, Chlorophyta) in India. Journal of Biosciences, 39(4), 711–716. https://doi.org/10.1007/s12038-014-9459-3

Bast, F., Bhushan, S., Rani, P., & John, A. A. (2016a). New record ofSargassum zhangii(Sargassaceae, Fucales) in India based on nuclear and mitochondrial DNA barcodes. Webbia, 71(2), 293–298. https://doi.org/10.1080/00837792.2016.1221188

Bast, F., John, A. A., & Bhushan, S. (2014a). Strong Endemism of Bloom-Forming Tubular Ulva in Indian West Coast, with Description of Ulva paschima Sp. Nov. (Ulvales, Chlorophyta). PLoS ONE, 9(10), e109295. https://doi.org/10.1371/journal.pone.0109295

Bast, F., John, A. A., & Bhushan, S. (2016b). Molecular assessment of invasive CarrageenophyteKappaphycus alvareziifrom India based on ITS-1 sequences. Webbia, 71(2), 287–292. https://doi.org/10.1080/00837792.2016.1221187

Bast, F., John, A. A., & Bhushan, S. (2015). Cladophora goensis sp. nov. (Cladophorales, Ulvophyceae) –a bloom forming marine algae from Goa, India. Ind J Geo-Mar Sci. 44(‘2): 1874–1879.

Belkacemi, L., Belalia, M., Djendara, A., & Bouhadda, Y. (2020). Antioxidant and antibacterial activities and identification of bioactive compounds of various extracts of Caulerpa racemosa from Algerian coast. Asian Pacific Journal of Tropical Biomedicine, 10(2), 87. https://doi.org/10.4103/2221-1691.275423

Beringuela, R. T., Purganan, D. J. E., Azanza, R. V., & Onda, D. F. L. (2020). Spatiotemporal variability and association of diatom-dinoflagellate assemblages of Acanthophora, Hypnea and Gracilaria (Rhodophyta). European Journal of Phycology, 55(3), 361–371. https://doi.org/10.1080/09670262.2020.1740797

Bermejo, R., Cara, C. L., Macías, M., Sánchez-García, J., & Hernández, I. (2020). Growth rates of Gracilariopsis longissima, Gracilaria bursa-pastoris and Chondracanthus teedei (Rhodophyta) cultured in ropes: implication for N biomitigation in Cadiz Bay (Southern Spain). Journal of Applied Phycology, 32(3), 1879–1891. https://doi.org/10.1007/s10811-020-02090-8

Bibi, R., Tariq, R. M., & Rasheed, M. (2020). Toxic assessment, growth disrupting and neurotoxic effects of red seaweeds’ botanicals against the dengue vector mosquito Aedes aegypti L. Ecotoxicology and Environmental Safety, 195, 110451. https://doi.org/10.1016/j.ecoenv.2020.110451

Blunt JW, B. R. Copp, W.-P. Hu, M. Munro, P. T. Northcote, M. R. Prinsep. (2007) Marine natural products. Nat. Prod. Rep., 24: 31–86.

Bonanno, G., Veneziano, V., & Piccione, V. (2020). The alga Ulva lactuca (Ulvaceae, Chlorophyta) as a bioindicator of trace element contamination along the coast of Sicily, Italy. Science of The Total Environment, 699, 134329. https://doi.org/10.1016/j.scitotenv.2019.134329

Bringloe, T. T., Saunders, G. W. DNA barcoding of the marine macroalgae from Nome, Alaska (Northern Bering Sea) reveals many trans-Arctic species. Polar Biol 42, 851–864 (2019). https://doi.org/10.1007/s00300-019-02478-4

Brodie, J., P.K. Hayes, G. L. Barker, L. M. Irvine and I. Bartsch. 1998. A reappraisal of Porphyra and Bangia (Bangiophycidae, Rhodophyta) in the Northeast Atlantic based on the rbcL–rbcS intergenic spacer. J. Phycol. 34: 1069–1074.

Butler, C. B., & Tankersley, R. A. (2020). Smells like home: The use of chemically-mediated rheotaxes by Limulus polyphemus larvae. Journal of Experimental Marine Biology and Ecology, 525, 151323. https://doi.org/10.1016/j.jembe.2020.151323

Dawes CJ, (1998) Marine Botany. John Wiley & Sons. pp. 1–459

De Clerck, O., M.D. Guiry, F. Leliaert, Y. Samyn and H. Verbruggen. 2013. Algal taxonomy: a road to nowhere? J. Phycol. 49: 215–225.

Dhargalkar V, Pereira N. (2005) Seaweed: Promising Plant of the Millennium. Science And Culture, 71(3-4), 60–66. http://drs.nio.org/drs/handle/2264/489

du Preez, R.; Paul, N.; Mouatt, P.; Majzoub, M. E.; Thomas, T.; Panchal, S. K.; Brown, L. Carrageenans from the Red Seaweed Sarconema filiforme Attenuate Symptoms of Diet-Induced Metabolic Syndrome in Rats. Mar. Drugs 2020, 18, 97.

El-Shaibany, A., AL-Habori, M., Al-Maqtari, T., & Al-Mahbashi, H. (2020). The Yemeni Brown Algae Dictyota dichotoma Exhibit High In Vitro Anticancer Activity Independent of Its Antioxidant Capability. BioMed Research International, 2020, 1–9. https://doi.org/10.1155/2020/2425693

El-Sheekh, M., El-Sabagh, S., Abou Elsoud, G. et al. Efficacy of Immobilized Biomass of the Seaweeds Ulva lactuca and Ulva fasciata for Cadmium Biosorption. Iran J Sci Technol Trans Sci 44, 37–49 (2020). https://doi.org/10.1007/s40995-020-00828-0

Ersalina, E. B., Abdillah, A. A., & Sulmartiwi, L. (2020). Potential of Caulerpa racemosa extracts as sunscreen creams. IOP Conference Series: Earth and Environmental Science, 441, 012007. https://doi.org/10.1088/1755-1315/441/1/012007

Fonseca, V. G., Carvalho, G. R., Sung, W., Johnson, H. F., Power, D. M., Neill, S. P., … Creer, S. (2010). Second-generation environmental sequencing unmasks marine metazoan biodiversity. Nature Communications, 1(1). https://doi.org/10.1038/ncomms1095

Ganesan, A. R., Subramani, K., Balasubramanian, B., Liu, W. C., Arasu, M. V., Al-Dhabi, N. A., & Duraipandiyan, V. (2020). Evaluation of in vivo sub-chronic and heavy metal toxicity of under-exploited seaweeds for food application. Journal of King Saud University - Science, 32(1), 1088–1095. https://doi.org/10.1016/j.jksus.2019.10.005

Ganesan, M., Trivedi, N., Gupta, V., Madhav, S. V., Radhakrishna Reddy, C., & Levine, I. A. (2019). Seaweed resources in India – current status of diversity and cultivation: prospects and challenges. Botanica Marina, 62(5), 463–482. https://doi.org/10.1515/bot-2018-0056

Ganesan, M.; Sahoo, N.; Eswaran, K. Raft culture of Gracilaria edulis in open sea along the south-eastern coast of India. Aquaculture 2011, 321, 145–151

Geoffroy, A., Le Gall, L., & Destombe, C. (2012). Cryptic introduction of the red alga Polysiphonia morrowii Harvey (Rhodomelaceae, Rhodophyta) in the North Atlantic Ocean highlighted by a DNA barcoding approach. Aquatic Botany, 100, 67–71. https://doi.org/10.1016/j.aquabot.2012.03.002

Govindaraju, K., Anand, K. V., Anbarasu, S., Theerthagiri, J., Revathy, S., Krupakar, P., Durai, G., Kannan, M., & Subramanian, K. S. (2020). Seaweed (Turbinaria ornata)-assisted green synthesis of magnesium hydroxide [Mg(OH)2] nanomaterials and their anti-mycobacterial activity. Materials Chemistry and Physics, 239, 122007. https://doi.org/10.1016/j.matchemphys.2019.122007

Graba-Landry, A. C., Loffler, Z., McClure, E. C., Pratchett, M. S., & Hoey, A. S. (2020). Impaired growth and survival of tropical macroalgae (Sargassum spp.) at elevated temperatures. Coral Reefs, 39(2), 475–486. https://doi.org/10.1007/s00338-020-01909-7

Granado, P., De Grande, F. R., & Costa, T. M. (2020). Association of Epialtus brasiliensis Dana, 1852 (Brachyura, Majoidea) with different species of seaweed. Nauplius, 28. https://doi.org/10.1590/2358-2936e2020004

Griffith, M. K., Schneider, C. W., Wolf, D. I., Saunders, G. W., & Lane, C. E. (2017). Genetic barcoding resolves the historically known red alga Champia parvula from southern New England, USA, as C. farlowii sp. nov. (Champiaceae, Rhodymeniales). Phytotaxa, 302(1), 77. https://doi.org/10.11646/phytotaxa.302.1.8

Gu, Z., Yang, L.-E., Chen, Z., & Chen, W. (2020). Comparative analysis of different DNA barcodes for applications in the identification and production of Pyropia. Algal Research, 47, 101874. doi.org/10.1016/j.algal.2020.101874

Guillemin, M.-L., González-Wevar, C., Cárdenas, L., Dubrasquet, H., Garrido, I., Montecinos, A., Ocaranza-Barrera, P., & Flores Robles, K. (2020). Comparative Phylogeography of Antarctic Seaweeds: Genetic Consequences of Historical Climatic Variations. In Antarctic Seaweeds (pp. 103–127). Springer International Publishing. https://doi.org/10.1007/978-3-030-39448-6_6

Hajibabaei, M., Shokralla, S., Zhou, X., Singer, G. A. C., & Baird, D. J. (2011). Environmental Barcoding: A Next-Generation Sequencing Approach for Biomonitoring Applications Using River Benthos. PLoS ONE, 6(4), e17497. https://doi.org/10.1371/journal.pone.0017497

Hamel, K., & Smith, C. M. (2020). Comparative time-courses of photoacclimation by Hawaiian native and invasive species of Gracilaria (Rhodophyta). Aquatic Botany, 163, 103210. https://doi.org/10.1016/j.aquabot.2020.103210

Harvey, A. S., Woelkerling, W. J., & de Reviers, B. (2020). A taxonomic analysis of Jania (Corallinaceae, Rhodophyta) in south-eastern Australia. Australian Systematic Botany. https://doi.org/10.1071/sb18064

Hebert, P. D. N., Cywinska, A., Ball, S. L. & deWaard, J. R. (2003) Biological identifications through DNA barcodes. Proceedings of the Royal Society B: Biological Sciences 270, 313–321, https://doi.org/10.1098/rspb.2002.2218

Heidari K, jafari D, babaei N, Esfandyari M. (2020) Removal of heavy metal zinc from aqueous solution by the alga Gracilaria corticata. IQBQ.; 4(1):51–43

Hleap, J. S., Littlefair, J. E., Steinke, D., Hebert, P. D. N., & Cristescu, M. E. (2020). Assessment of current taxonomic assignment strategies for metabarcoding eukaryotes. Cold Spring Harbor Laboratory. https://doi.org/10.1101/2020.07.21.214270

Hollingsworth PM, Forrest LL, Spouge JL, Hajibabaei M, Ratnasingham S et al. (2009) A DNA barcode for land plants. Proc Natl Acad Sci U S A 106: 12794–12797. https://doi:10.1073/pnas.0905845106

Imchen T and Ezaz W. (2018) Time course nutrient uptake study of some intertidal rocky shore macroalgae and the limiting effect due to synergistic interaction. Indian Journal of Geo Marine Sciences. 49(02): 287–292.

Jha, B., Reddy, C. R. K., Thakur, M. C., & Rao, M. U. (2009). Seaweeds of India. Springer Netherlands. https://doi.org/10.1007/978-90-481-2488-6

Kambey, C. S. B., Sondak, C. F. A., & Chung, I. (2020). Potential growth and nutrient removal of Kappaphycus alvarezii in a fish floating net cage system in Sekotong Bay, Lombok, Indonesia. Journal of the World Aquaculture Society. https://doi.org/10.1111/jwas.12683

Kannan, L., & Thangaradjou, T. (2007). PROSPECTS OF SEAWEED CULTIVATION IN INDIA VIS-A-VIS WORLD. Acta Horticulture, 742, 191–195. https://doi.org/10.17660/actahortic.2007.742.25

Kayombo, C. J., Barnabas, G., & Likinguraine, K. (2020). Evaluation of Seagrass and Seaweed Species Diversity, Abundance, and Human Activities Endangering their Existence at Indian Ocean Shore at Kigamboni in Dar es Salaam City, Coastal Zone of Tanzania. East African Journal of Environment and Natural Resources, 2(2), 19–26. https://doi.org/10.37284/eajenr.2.2.174

Kazi MA, Reddy CRK, Jha B (2013) Molecular Phylogeny and Barcoding of Caulerpa (Bryopsidales) Based on the tufA, rbcL, 18S rDNA and ITS rDNA Genes. PLoS ONE 8(12): e82438. https://doi:10.1371/journal.pone.0082438

Kh. Gabr, M., F. Ziena, A., & M. Hellal, A. (2020). Abundance and diversity of amphipod species associated with macro-algae at Ras-Mohamed, Aqaba Gulf, Red Sea, Egypt. Egyptian Journal of Aquatic Biology and Fisheries, 24(3), 1–15. https://doi.org/10.21608/ejabf.2020.103630

Khan SI., and Satam S., (2003) Seaweed mariculture: scope and potential in India. Aquacult. Asia., 8: 26–29

Kimura M. (1980). A simple method for estimating evolutionary rate of base substitutions through comparative studies of nucleotide sequences. Journal of Molecular Evolution 16:111–120.

Krishnamoorthy S, V. Venkatesalu, G. A. R. M. C. (2015). Antibacterial Activity of Diff Erent Solvent Extracts of Caulerpa Chemnitzia (Esper) J.V. Lamououx, from Mandapam, Gulf of Mannar Southeast Coast, Tamil Nadu, India”. Journal of Medicinal Herbs and Ethnomedicine. 1: 24–31.

Kucera, H., & Saunders, G. W. (2008). Assigning morphological variants of Fucus (Fucales, Phaeophyceae) in Canadian waters to recognized species using DNA barcoding. Botany, 86(9), 1065–1079. https://doi.org/10.1139/b08-056

Kumar, A., Kumar, J., & Bhaskar, T. (2020). High surface area biochar from Sargassum tenerrimum as potential catalyst support for selective phenol hydrogenation. Environmental Research, 186, 109533. https://doi.org/10.1016/j.envres.2020.109533

Lanham, B. S., P. E. Gribben, and A. G. Poore. 2015. Beyond the border: effects of an expanding algal habitat on the fauna of neighboring habitats. Marine Environmental Research 106:10–18.

Lavaud, R., Filgueira, R., Nadeau, A., Steeves, L., & Guyondet, T. (2020). A Dynamic Energy Budget model for the macroalga Ulva lactuca. Ecological Modelling, 418, 108922. https://doi.org/10.1016/j.ecolmodel.2019.108922

Lee, H. W., & Kim, M. S. (2015). Species delimitation in the green algal genus Codium (Bryopsidales) from Korea using DNA barcoding. Acta Oceanologica Sinica, 34(4), 114–124. https://doi.org/10.1007/s13131-015-0651-6

Leray, M., & Knowlton, N. (2015). DNA barcoding and metabarcoding of standardized samples reveal patterns of marine benthic diversity. Proceedings of the National Academy of Sciences, 112(7), 2076–2081. https://doi.org/10.1073/pnas.1424997112

Lula Leite, D. S., Tavares Pessoa Pinho de Vasconcelos, E. R., Riul, P., Alves de Freitas, N. D., & Cavalcanti de Miranda, G. E. (2020). Evaluation of the conservation status and monitoring proposal for the coastal reefs of Paraíba, Brazil: Bioindication as an environmental management tool. Ocean & Coastal Management, 194, 105208. https://doi.org/10.1016/j.ocecoaman.2020.105208

Manikantan, G., PrasannaKumar, C., Vijaylaxmi, J., Pugazhvendan, S. R., & Prasanthi, N. (2020). Diversity, phylogeny, and DNA barcoding of brachyuran crabs in artificially created mangrove environments. Cold Spring Harbor Laboratory. https://doi.org/10.1101/2020.09.07.286823

Mahendran S and Saravanan S (2014) Molecular taxonomy of green seaweeds Ulva lactuca and Caulerpa taxifolia through phylogenetic analysis. Ind J of Geo Mar Sci. 46(2):414–419.

Mantri, V. A., Kavale, M. G., & Kazi, M. A. (2019). Seaweed Biodiversity of India: Reviewing Current Knowledge to Identify Gaps, Challenges, and Opportunities. Diversity, 12(1), 13. https://doi.org/10.3390/d12010013

Méndez, F., Marambio, J., Ojeda, J., Rosenfeld, S., Rodríguez, J., Tala, F., & Mansilla, A. (2019). Variation of the photosynthetic activity and pigment composition in two morphotypes of Durvillaea antarctica (phaeophyceae) in the sub-Antarctic ecoregion of Magallanes, Chile. Journal of Applied Phycology, 31, 905–913. https://doi.org/10.1007/s10811-018-1675-z

Mineur, F., Le Roux, A., Stegenga, H. et al. Four new exotic red seaweeds on European shores. Biol Invasions 14, 1635–1641 (2012). https://doi.org/10.1007/s10530-012-0186-0

Mitchell, A. (2008) DNA barcoding demystified. Australian Journal of Entomology 47, 169–173, https://doi.org/10.1111/j.1440-6055.2008.00645.x

Montes, M., Rico, J. M., García-Vazquez, E., & Borrell Pichs, Y. J. (2017). Molecular barcoding confirms the presence of exotic Asian seaweeds (Pachymeniopsis gargiuliandGrateloupia turuturu) in the Cantabrian Sea, Bay of Biscay. PeerJ, 5, e3116. https://doi.org/10.7717/peerj.3116

Murugaiyan K (2020) Phytochemical composition on three species of Sargassum from southeast coast of Tamil Nadu, India. Plant Archives, 20(1): 1555–1559.

Murugaiyan K and Sivakumar K (2020) Seasonal variation in biochemical and elemental composition of some marine algae of Mandapam, Southeast coast of India. Plant Archives, 20(1): 1987–1993.

Nasab, S. B., Homaei, A., & Karami, L. (2020). Kinetic of α-amylase inhibition by Gracilaria corticata and Sargassum angustifolium extracts and zinc oxide nanoparticles. Biocatalysis and Agricultural Biotechnology, 23, 101478. https://doi.org/10.1016/j.bcab.2019.101478

Nei M. and Kumar S. (2000). Molecular Evolution and Phylogenetics. Oxford University Press, New York.

Nguyen, T. V., & Boo, S. M. (2020). Distribution patterns and biogeography of Sargassum (Fucales, Phaeophyceae) along the coast of Vietnam. Botanica Marina, 0(0). https://doi.org/10.1515/bot-2019-0082

Nogueira, M. C. F., Henriques, M. B. Large-scale versus family-sized system production: economic feasibility of cultivating Kappaphycus alvarezii along the southeastern coast of Brazil. J Appl Phycol 32, 1893–1905 (2020). https://doi.org/10.1007/s10811-020-02107-2

Osman, M. E. H., Abo-Shady, A. M., Elshobary, M. E., Abd El-Ghafar, M. O., & Abomohra, A. E.-F. (2020). Screening of seaweeds for sustainable biofuel recovery through sequential biodiesel and bioethanol production. Environmental Science and Pollution Research. https://doi.org/10.1007/s11356-020-09534-1

O’Sullivan L, B. Murphy, P. McLoughlin, P. Duggan, P. G. Lawlor, H. Hughes, G. E., (2010) Gardiner Prebiotics from marine macroalgae for human and animal health applications. Mar. Drugs, 8: 2038–2064

Pasqualetti, M., Giovannini, V., Barghini, P., Gorrasi, S., & Fenice, M. (2020). Diversity and ecology of culturable marine fungi associated with Posidonia oceanica leaves and their epiphytic algae Dictyota dichotoma and Sphaerococcus coronopifolius. Fungal Ecology, 44, 100906. https://doi.org/10.1016/j.funeco.2019.100906

Pooja, K., Rani, S., Rana, V., & Pal, G. K. (2020). Aquatic plants as a natural source of antimicrobial and functional ingredients. In Functional and Preservative Properties of Phytochemicals (pp. 93–118). Elsevier. https://doi.org/10.1016/b978-0-12-818593-3.00003-8

Porzio, L., Arena, C., Lorenti, M., De Maio, A., & Buia, M. C. (2020). Long-term response of Dictyota dichotoma var. intricata (C. Agardh) Greville (Phaeophyceae) to ocean acidification: Insights from high pCO2 vents. Science of The Total Environment, 731, 138896. https://doi.org/10.1016/j.scitotenv.2020.138896

Prasannakumar, C., Manikantan, G., Vijaylaxmi, J., Gunalan, B., Manokaran, S., & Pugazhvendan, S. R. (2020). Strengthening of marine amphipod DNA barcode libraries for environmental monitoring. Cold Spring Harbor Laboratory. https://doi.org/10.1101/2020.08.26.268896

Rajaram, R., Rameshkumar, S., & Anandkumar, A. (2020). Health risk assessment and potentiality of green seaweeds on bioaccumulation of trace elements along the Palk Bay coast, Southeastern India. Marine Pollution Bulletin, 154, 111069. https://doi.org/10.1016/j.marpolbul.2020.111069

Rajivgandhi, G. N., Ramachandran, G., Maruthupandy, M., Manoharan, N., Alharbi, N. S., Kadaikunnan, S., Khaled, J. M., Almanaa, T. N., & Li, W.-J. (2020). Anti-oxidant, antibacterial and anti-biofilm activity of biosynthesized silver nanoparticles using Gracilaria corticata against biofilm producing K. pneumoniae. Colloids and Surfaces A: Physicochemical and Engineering Aspects, 600, 124830. https://doi.org/10.1016/j.colsurfa.2020.124830

Rao, M. U.; Chaugule, B. B. Endangered and extinct seaweeds of Indian shore. In Recent Advances on Applied Aspects of Indian Marine Algae with Reference to Global Scenario; Tewari, A., Ed.; Central Salt and Marine Chemicals Research Institute: Bhavnagar, India, 2006; pp. 141–146.

Rashad, S., & A. El-Chaghaby, G. (2020). Marine Algae in Egypt: distribution, phytochemical composition and biological uses as bioactive resources (a review). Egyptian Journal of Aquatic Biology and Fisheries, 24(5), 147–160. https://doi.org/10.21608/eiabf2020.103630

Renuka, R. R., Ravindranath, R. R. S., Raguraman, V. et al. In Vivo Toxicity Assessment of Laminarin Based Silver Nanoparticles from Turbinaria ornata in Adult Zebrafish (Danio rerio). J Clust Sci 31, 185–195 (2020). https://doi.org/10.1007/s10876-019-01632-6

Ribeiro, N. A., Chaves, H. V., da Conceição Rivanor, R. L., do Val, D. R., de Assis, E. L., Silveira, F. D., Gomes, F. I. F., Freitas, H. C., Vieira, L. V., da Silva Costa, D. V., de Castro Brito, G. A., Bezerra, M. M., & Benevides, N. M. B. (2020). Sulfated polysaccharide from the green marine algae Caulerpa racemosa reduces experimental pain in the rat temporomandibular joint. International Journal of Biological Macromolecules, 150, 253–260. https://doi.org/10.1016/j.ijbiomac.2020.01.272

Rodríguez, C. Y., & Otaíza, R. D. (2020). Morphological variability in a red seaweed: confirmation of co□occurring f. lessonii and f. chauvinii in Chondracanthus chamissoi (Rhodophyta, Gigartinales). Journal of Phycology, 56(2), 469–480. https://doi.org/10.1111/jpy.12955

Rushdi, M. I., Abdel-Rahman, I. A. M., Saber, H., Attia, E. Z., Abdelraheem, W. M., Madkour, H. A., & Abdelmohsen, U. R. (2020). The genus Turbinaria: chemical and pharmacological diversity. Natural Product Research, 1–19. https://doi.org/10.1080/14786419.2020.1731741

Rushdi, M. I., Abdel-Rahman, I. A. M., Saber, H., Attia, E. Z., Abdelraheem, W. M., Madkour, H. A., Hassan, H. M., Elmaidomy, A. H., & Abdelmohsen, U. R. (2020). Pharmacological and natural products diversity of the brown algae genus Sargassum. RSC Advances, 10(42), 24951–24972. https://doi.org/10.1039/d0ra03576a

Sanders ER, Karol KG, McCourt RM (2003) Occurrence of matK in a trnK group II intron in Charophyte green algae and phylogeny of the Characeae. Am J Bot 90:628–633. https://doi:10.3732/ajb.90.4.628

Sathakit, R., Ruangchuay, R., Luangthuvapranit, C., & Bovornruangroj, N. (2020). Effect of Carbon dioxide (CO2) concentration on Growth of Ulva intestinalis in Photobioreactor. IOP Conference Series: Earth and Environmental Science, 416, 012020. https://doi.org/10.1088/1755-1315/416/1/012020

Saunders, G. W. (2009). Routine DNA barcoding of Canadian Gracilariales (Rhodophyta) reveals the invasive speciesGracilaria vermiculophyllain British Columbia. Molecular Ecology Resources, 9, 140–150. https://doi.org/10.1111/j.1755-0998.2009.02639.x

Saunders, G. W., & McDevit, D. C. (2013). DNA barcoding unmasks overlooked diversity improving knowledge on the composition and origins of the Churchill algal flora. BMC Ecology, 13(1), 9. https://doi.org/10.1186/1472-6785-13-9

Saunders, G. W., & Moore, T. E. (2013). Refinements for the amplification and sequencing of red algal DNA barcode and RedToL phylogenetic markers: a summary of current primers, profiles and strategies. ALGAE, 28(1), 31–43. https://doi.org/10.4490/algae.2013.28.1.031

Schaffelke, B., J.E. Smith and C.L. Hewitt. 2006. Introduced macroalgae – a growing concern. J. Appl. Phycol. 18: 529–541.

Siddiqui Z.H., Abbas Z.K., Hakeem K.R., Khan M.A., Ilah M.A. (2020) A Molecular Assessment of Red Algae with Reference to the Utility of DNA Barcoding. In: Trivedi S., Rehman H., Saggu S., Panneerselvam C., Ghosh S. (eds) DNA Barcoding and Molecular Phylogeny. Springer, Cham. https://doi.org/10.1007/978-3-030-50075-7_7

Sirison, N., & Burnett, N. P. (2019). Turbinaria ornata (Phaeophyceae) varies size and strength to maintain environmental safety factor across flow regimes. Journal of Phycology, 56(1), 233–237. https://doi.org/10.1111/jpy.12933

Starko, S., Soto Gomez, M., Darby, H., Demes, K. W., Kawai, H., Yotsukura, N., … Martone, P. T. (2019). A comprehensive kelp phylogeny sheds light on the evolution of an ecosystem. Molecular Phylogenetics and Evolution, 136, 138–150. https://doi.org/10.1016/j.ympev.2019.04.012

Stelling□Wood, T. P., Gribben, P. E., & Poore, A. G. B. (2020). Habitat variability in an underwater forest: Using a trait based approach to predict associated communities. Functional Ecology, 34(4), 888–898. https://doi.org/10.1111/1365-2435.13523

Subramaniam, D., Hanna, L. E., Maheshkumar, K., Ponmurugan, K., Al-Dhabi, N. A., & Murugan, P. (2020). Immune stimulatory and anti-HIV-1 potential of extracts derived from marine brown algae Padina tetrastromatica. Journal of Complementary and Integrative Medicine, 17(2). https://doi.org/10.1515/jcim-2019-0071

Subramanian G, Nagaraj A, Sasikala J, Ravi P and Sona P (2020) Nutrient content of red algal species of a genus Gracilaria from the coastal areas of Rameswaram, Tamil Nadu, India. The Int. J. of Anal. And Exp. Mod. Anal. 12(5): 1721–1727.

Susanto, E.; Fahmi, A. S.; Hosokawa, M.; Miyashita, K. Variation in lipid components from 15 species of tropical and temperate seaweeds. Mar. Drugs 2019, 17, 630.

Tajima F. (1989). Statistical methods to test for nucleotide mutation hypothesis by DNA polymorphism. Genetics 123:585–595.

Tan J, Lim P-E, Phang S-M, Hong DD, Sunarpi H, et al. (2012) Assessment of Four Molecular Markers as Potential DNA Barcodes for Red Algae Kappaphycus Doty and Eucheuma J. Agardh (Solieriaceae, Rhodophyta). PLoS ONE 7(12): e52905. https://doi:10.1371/journal.pone.0052905

Tan, T.-T., Song, S.-L., Poong, S.-W., Ward, G. M., Brodie, J., & Lim, P.-E. (2020). The effect of grazing on the microbiome of two commercially important agarophytes, Gracilaria firma and G. salicornia (Gracilariaceae, Rhodophyta). Journal of Applied Phycology. https://doi.org/10.1007/s10811-020-02062-y

Tanaka, Y., Ashaari, A., Mohamad, F. S., & Lamit, N. (2020). Bioremediation potential of tropical seaweeds in aquaculture: low-salinity tolerance, phosphorus content, and production of UV-absorbing compounds. Aquaculture, 518, 734853. https://doi.org/10.1016/j.aquaculture.2019.734853

Thankaraj, Sekar S R, V., Kumaradhass, H. G., Perumal, N., & Hudson, A. S. (2019). Exploring the antimicrobial properties of seaweeds against Plasmopara viticola (Berk. and M.A. Curtis) Berl. and De Toni and Uncinula necator (Schwein) Burrill causing downy mildew and powdery mildew of grapes. Indian Phytopathology, 73(2), 185–201. https://doi.org/10.1007/s42360-019-00137-6

Tukarambai, M., & Venakateswarlu, P. (2020). A study of lead removal using sargassum tenerrimum (brown algae): Biosorption in column study. Materials Today: Proceedings, 27, 421–425. https://doi.org/10.1016/j.matpr.2019.11.254

Vasuki S, Kokilam G, and Babitha D (2020) Mineral composition of some selected brown seaweeds from Mandapam region of Gulf of Mannar, Tamil Nadu. Indian Journal of Geo Marine Sciences. 49(01), 63–66.

